# *In silico* design of DNA sequences for *in vivo* nucleosome positioning

**DOI:** 10.1101/2023.05.15.540782

**Authors:** Ethienne Routhier, Edgar Pierre, Alexandra Joubert, Astrid Lancrey, Jean-Baptiste Boulé, Julien Mozziconacci

## Abstract

The computational design of synthetic DNA sequences with desired in vivo properties is gaining traction in the field of synthetic genomics. We propose here a computational method which combines a kinetic Monte Carlo framework with a deep mutational screening based on deep learning predictions. We apply our method to build regular nucleosome arrays with tailored nucleosomal repeat lengths (NRL) in yeast. Our design is validated in vivo by successfully engineering and integrating thousands of kilobases long tandem arrays of computationally optimized sequences which could accommodate NRLs much larger than the yeast natural NRL. This method delineates the key sequence rules for nucleosome positioning in yeast and is readily applicable to other sequence properties and other genomes.

## 1 Introduction

Recent biotechnology techniques such as CRISPR/Cas9 and DNA oligonucleotide *in vivo* assembly have opened ways to precisely and extensively modify genomes. Taking advantage of these technologies, several projects have been launched with the aim to partially or completely design and assemble synthetic genomes [1]. Nevertheless, controlling chromatin assembly and gene expression on a synthetic genome remains a challenge. Decisive efforts have been recently made for designing promoter sequences that produce controlled levels of mRNA in yeast [2, 3] or the activity of enhancers in a Drosophila cell line [4]. While the control of gene expression is now within our grasp, there is yet no efficient way to control by sequence design the positioning of nucleosomes along a synthetic DNA cassette inserted in an eukaryotic genome. Nucleosome positioning is however of crucial importance as it influences DNA accessibility to DNA binding factors involved in DNA replication and transcription, thus adding a supplementary level of control on top of the DNA sequence [5, 6]. In a seminal experiment, Lowary and Widom used a SELEX approach to isolate a DNA sequence with the highest affinity to a histone octamer among a set of more than 5 × 10^12^ sequences [7]. The resulting 147 bp sequence, known as the 601-sequence, has been extensively used to reconstruct regular nucleosomal arrays *in vitro* (e.g. [8]). Given the success in using the 601 sequence for nucleosome positioning in *vitro*, we recently addressed the affinity of nucleosomes to the 601 sequence *in vivo* using insertions in the *S. cerevisiae* genome of three long arrays of approximately 50 nucleosomes, with three different spacing between nucleosomes (167,197 and 237 bp), also known as nucleosomal repeat length (NRL). In this study we found that in sharp contrast with the *in vitro* experiments, the affinity of nucleosomes to the 601 sequence was very low *in vivo* [9] suggesting that this sequence is not an adequate tool to control nucleosome positions in synthetic genomics approaches. Computational tools are a good alternative to optimize the design of synthetic sequences from *in vivo* data. Among the available computational methods, deep learning has been widely applied to building predictive models that relate DNA sequences to genomic functions [10, 11]. The ability of deep neural networks to predict annotations resulting from variations of a sequence is now used for *de novo* design of genomic sequences, including tailored alternative poly-adenylation sites [12, 13] or human 5’UTR sequences [14, 15].

Building on these previous studies, we use here a nucleosome density predictor [16] together with a kinetic Monte-Carlo framework in order to design three sequences (of 167 bp, 197 bp and 237 bp) that lead to a regular nucleosome positioning when assembled into tandem repeated arrays *in vivo* (Fig.1). The three NRL chosen encompassed the natural NRL of *S*.*cerevisiae* (167 bp) and the longest NRL known in eukaryotic species (237 bp, found in the sea urchin sperm [17]).

**Figure 1:**
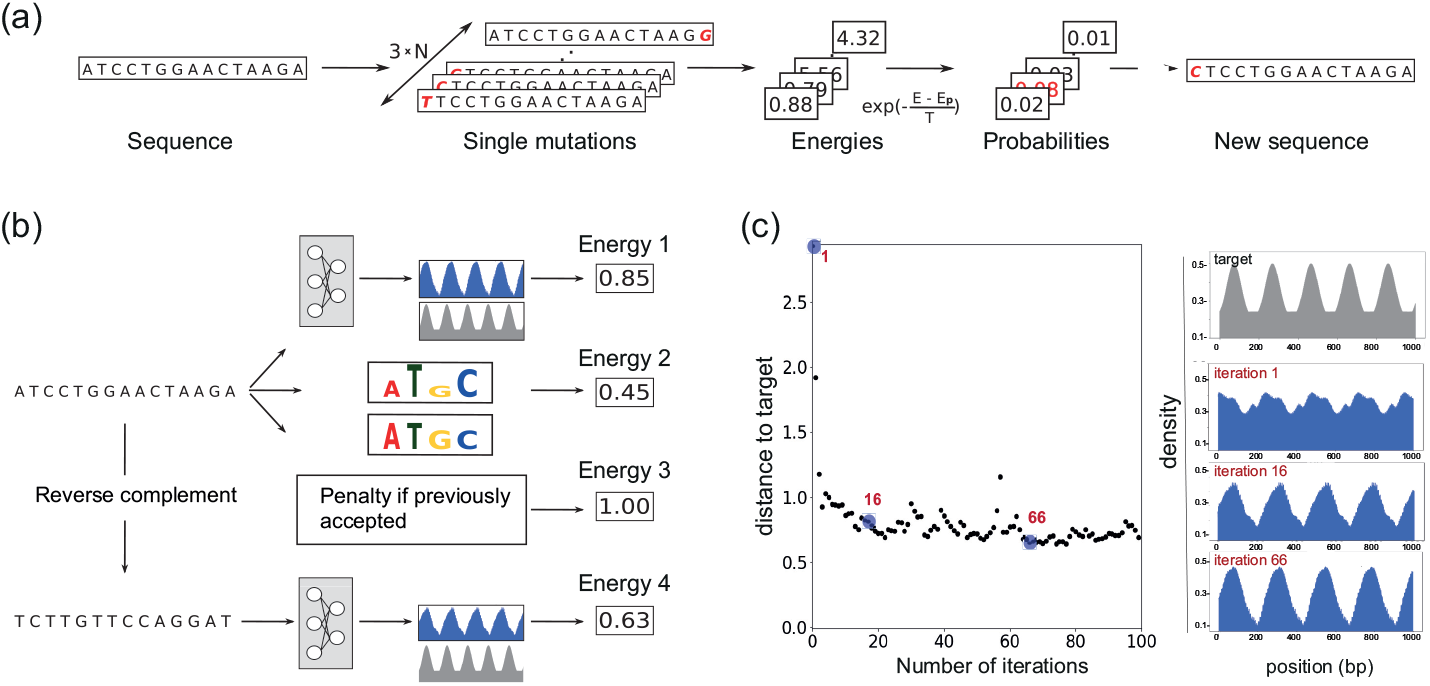
Sequence optimization via kinetic Monte-Carlo. (a) General principle of the method. (b) The energy associated to a sequence is the sum of four terms: (1) the distance between the nucleosome density predicted by the model and the target nucleosome density (2) the distance between the GC content of the sequence and the natural GC content of the yeast DNA (3) a penalty if the sequence has been already sampled (4) the distance between the density predicted on the reversed complemented sequence and the target density. (c) Evolution of the predicted nucleosome density (blue densities on the right) and of the corresponding energy (surrounded in blue) during the optimization process. The target density is displayed at the top (grey).

## 2 MATERIALS AND METHODS

### Synthetic sequence initialization

The initial sequences were determined at random. Each of the four (A,T,G,C) nucleotides is sequentially picked *N* times with a probability proportional to its abundance within the genome. This procedure insures that starting sequences have a GC content similar to the natural GC content of *S*.*cerevisiae*.

### Synthetic sequence mutation

Our mutation and selection strategy for optimizing the sequence is inspired from the kinetic Monte-Carlo (k-MC) method originally designed for Ising spin systems ([19]). Once the sequence of length *N* is initialized, it is duplicated 3 × *N* times in order to create 3 × *N* new sequences which will each harbor one of the 3 × *N* single mutations of the sequence (Fig.1a). We then associate an energy term with each of the mutated sequences (see below). For every sequence *i* a selection probability Γ_*i*_ is then defined as follow:

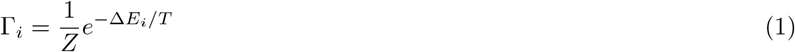

with *Z* being the normalisation factor 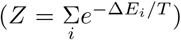 and *T* the temperature, a broadening factor chosen by the user and corresponding to a classical temperature in a Maxwell-Boltzmann distribution. The next sequence is then randomly chosen according to this distribution of probability. The *i*^*th*^ configuration is selected - with a probability Γ_*i*_ - and the process continues for a given number of steps (10^4^ in our case). Each chosen configuration is saved at each step, and the configuration with the minimal energy is chosen at the end.

### Synthetic sequence energy term

An energy is associated to every sequence, representing how far the sequence is from having the desired characteristics. The energy can be divided into four parts (Fig.1 b), described below.

#### Nucleosome density energy on the direct strand E_reg_

In a former work we used a convolutional neural network (CNN) to predict the nucleosome density at the center nucleotide of a 2001 bp long DNA sequence [16]. In order to evaluate the nucleosome density over our synthetic array, we first create a 2001 + *N* bp long sequence by repeating the monomeric synthetic sequence. Then, this sequence is cut in *N* sequences of 2001 bp long used as inputs of the network thus providing the predicted nucleosome density over the whole synthetic sequence. The nucleosome density related energy is the distance between the predicted nucleosome density on the synthetic sequence and a target density. It forces the predicted density to converge to the target density (Fig.1 c) The distance is defined as :

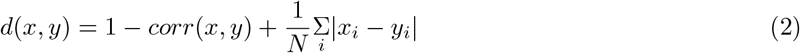

where corr stands for the Pearson correlation and *N* is the length of the sequence. The target signal *y*^*target*^ is a combination of a gaussian distribution in the nucleosome region and a uniform distribution in the linker region:

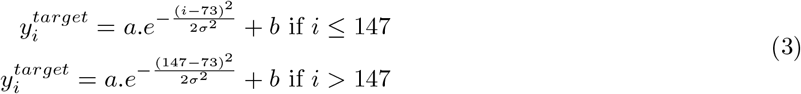

The parameters *a, b* and *σ* are chosen in order to create a realistic target (i.e which shape is similar to canonical nucleosomal densities found on the genome). The results are shown for *a* = 0.4, *b* = 0.2 and 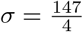.

#### GC content related energy E_GC_

As said previously, there are 4^*N*^ different sequences of a given length *N*. The CNN learned to position nucleosomes for sequences within the genome of *saccharomyces cerevisiae*. As a result, one wants to get a synthetic sequence which has a rather similar GC content as natural yeast. In order to do so, a first constrain energy is calculated as follow:

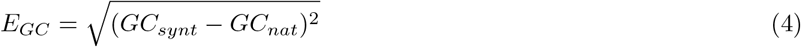

with *GC*_*synt*_ the GC content of the synthetic sequence and *GC*_*nat*_ the GC content of the natural yeast (in that case, *GC*_*nat*_ ≈ 0.38).

#### Enhanced sampling mutation energy E_mut_

It is common that such monte carlo procedure can get stuck into a local minimum of the energy, for instance flipping between the same two mutations. To avoid this potential pitfall, we added a term to the energy term that penalizes sequences that were already generated during the optimization process.

#### Nucleosome density energy on the reverse strand E_rev_

This energy is calculated in the same way as the nucleosome density related energy on the direct strand, but using the reverse complement strand instead. On most sequences this term is not relevant since the nucleosomal density predicted by the network is, due to the training process, the same for both strands. However, we added this term to add an extra penalty to sequences for which the network would not work as expected and predict different densities for the two strands.

As a result, Eq.1 can be rewritten as:

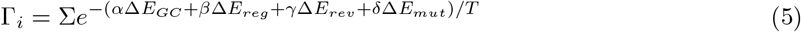

with *α, β, γ* and *δ* the relative weights for each energy component, chosen to get the same order of magnitude (e.g. = 1) for each energy term.

### Interpretation of the mutations selected during the optimization process

#### Positions of the mutations along the sequence

A thousand independent k-MC optimization processes were executed and stopped after 20 steps. At each time step, the energy associated with each possible mutation is computed and stored. To quantify the importance of a given nucleotide in the positioning capacity of the sequence, the average absolute energy variation created by a mutation at this specific position is computed.

#### Typical mutation logos

For each length (167, 197 and 237 bp), we collected all the 5-bp long motifs centered at each mutation site during 1000 independent optimization processes stopped after 20 steps. We then selected motifs for which a mutation induces an absolute energy variation higher than 0.5 (see **Supplementary Figure S1** for the distribution of the absolute energy changes). We then regrouped the collected motifs into four categories depending on the sign of the energy modification and on their position, located in the linker region or in the dyad region (respectively after 147 bp and between 50 bp and 100 bp). We ended up with 14938 negative mutations in the linker, 16165 negative mutations in the dyad, 52603 positives mutations in the dyad and 158422 positive mutations in the linker. For each of these four categories we extracted significant logos of 5 bp using STREME [20, 18] with the argument w=5. We also provided a list of control sequence to STREME by passing the list of 5 bp motifs associated to a mutation that modify the energy by less than 0.03. Finally, we selected logos that matched more than 10 % of the input sequences and represented them before and after the mutation.

### In vivo assembly of the computationally designed sequences

#### Strains, plasmids, reagents and media

In vivo genomic assembly of synthetic sequences nucleosome positioning (SSNP, Supplementary Table 1) DNA repeats was performed in the yeast strain YPH499. Strains derived from this study are listed in Supplementary Table 2. Plasmids used for the *in vivo* expression of spCas9 and the guide RNA targeting the YMR262 gene have been previously described [21] and are listed in Supplementary Table 3. Strains were grown at 30°C in yeast extract Peptone Dextrose 2% media (YPD) or in the appropriate synthetic complete Dextrose 2% media (SCD), media minus relevant amino acids necessary to maintain plasmid borne auxotrophic markers. All media reagents were purchased from Formedium and used as recommended. Oligonucleotides used in this study were synthetized by Eurogentec. Enzymes for nucleic acids modification were purchased from New England Biolabs. Zymolyase 20T was purchased from Amsbio.

#### Assembly of synthetic sequences nucleosome positioning DNA repeats in the yeast genome

*In vivo* assembly using CRISPR/Cas9 and overlapping oligonucleotides in the Chromosome XIII of *S. cerevisiae* strain YPH499 was performed as previously described [21, 9]. In summary, donor DNA containing the left (YMR/SSNP) and the right (SSNP/YMR) genomic junction were amplified by PCR using primers couples O-1/O-2(O-3/O-4/O-5) and O-10/O-6(O-7/O-8/O-9) respectively. This donor DNA results, upon recombinational assembly in the deletion of the region -129 to 232 bp of the YMR gene. All oligonucleotides used for *in vivo* repeat assembly are listed in Supplementary Table 4. YPH499 was transformed with the Cas9 expressing plasmid pRS413-Cas9-His (AJ-P1). The resulting strain was transformed using the LiAc technique [22] with 1 μg of gRNA expressing plasmid targeting YMR262 (AJ-P2), 100 pmol of each of the four or six appropriate SSNP-oligonucleotides, and 10 pmol of both YMR/SSNP left and right junction PCR. After transformation cells were plated on SCD-His-Ura to select cells carrying both CAS9 and gRNA expressing plasmids. The in vivo assembly of synthetic sequences repeated arrays was verified by left and right junction PCR amplification (O-1/O-2 ; O-3 ; O-4 ; O-5 and O-10/O-6 ; O-7 ; O-8 ; O-9) and Sanger sequencing. The correct locus and the size of the SSNP assembly was confirm by analyzed recombinant clones by southern-blotting. Genomic DNA from recombinant strains were digested with BamHI and DraI, which cut at each side of the insertion locus. Digested DNA was electrophoresed in 1% agarose and transferred by capillarity onto a nylon membrane (Hybond N+, GE healthcare). Membranes were hybridized in Church buffer at 68°C with Cy5-probe ([23]). The genomic probe was a 1 kb DNA fragment amplified by PCR from genomic DNA using primers O-33/O-34 and Cy5-dCTP. Membranes were scanned using a FLA 9500 GE healthcare.

#### Mononucleosome preparation using Micrococcal Nuclease digestion and sequencing (Mnase-Seq)

Each strain was grown to an OD600 of 0.8 in 250 mL SCD media at 30°C with 200 rpm shaking. Cultures were treated with a final concentration of 1.85 % formaldehyde for 30 min at 30°C. Cross-linking was stopped by addition of 105 mM Glycine (final concentration). Cell pellets (6500g, 10 min) were washed and resuspended in 50 mL of 1 M Sorbitol, 10 mM Tris pH 7.5 supplemented with 10 mM ß-mercaptoethanol and 15 mg of Zymolyase 20T, and incubated 1h at 30°C with 50 rpm shaking. Spheroplasts were pelleted (6500g, 10 min), and resuspended in 2.4 mL solution containing 1M Sorbitol, 50 mM NaCl, 10 mM Tris pH 7.5, 5 mM MgCl2, 1 mM CaCl2 and 0.75 % Igepal CA630 freshly supplemented with 1 mM ß-mercaptoethanol, 500 μM spermidine and 1000 units of MNase. The spheroplasts/MNase mixture was then incubated at 37°C for 30 min and stopped by adding 600 μL of 1% SDS, 10 mM EDTA. Reversal of crosslink and protein removal was achieved by adding 0.6 mg of Proteinase K (ThermoFisher) and overnight incubation at 65°C. DNA was purifed using phenol/chloroform precipitation method. Mononucleosomal DNA was isolated on a 1.8 % agarose electrophoresis gel and purified using QIAquick Gel extraction Kit. After DNA concentration determination using a Qubit Fluorometer, samples were treated and sequenced on a NovaSeq 6000 S4 PE150 XP by the Eurofins sequencing platform (NGSelect Amplicons).

#### Construction of synthetic reference genomes

For the three synthetic sequences we constructed a reference genome carrying two tandem repeats plus 65 bp of a third one. Indeed, the construction of the reference genome is required to take into account all possible alignments of a reads. Reads for which one mate overlaps the linker and the beginning of the following synthetic sequence justify the use of a second repeat in our reference genome. Extreme cases where the 2 mates of a read overlap two linkers justify the use of the beginning on a third repeat (65 bp is the length of our read -1 bp).

#### Reads alignment

After the removal of barcodes, paired-end reads of 66 nt each were mapped against the appropriate (167, 197 or 237 bp) reference genome using Bowtie 2 (version 2.3.1) [24, 25] allowing at most 2 alignments per read (-k m) and fragments length ranging 100 to 200 bp for 167 and 197 strains, and 120 to 250 bp for 237 strains (-I and -X parameters). Concordant reads were specifically selected using samtools (version 1.10).

#### Reconstruction of sequence coverage over DNA repeats

By choosing a reference genome with two repeats and 65 bp of an additional repeat, a large proportion of the reads of interest should be able to map twice justifying the allowing of 2 hits per reads using bowtie 2. Reads presenting multiple hits were filtered to select the most 5’-end alignment. Then, the density of nucleosome (*i*.*e* DNA fragments) on the specific reference genome is reconstructed using bedtools (version 2.29.2). Finally, the density on a unique repetition of a synthetic sequence was calculated by adding the density on every repeats in the synthetic reference genome.

## 3 RESULTS AND DISCUSSION

### 3.1 Our computational model predicts the inefficiency of the Widom-601 sequence *in vivo*

In order to test the *in vivo* ability of the Widom-601 sequence to form regular nucleosome arrays of various NRLs we recently synthesize and integrated such arrays within a yeast chromosome [9]. Our results unambiguously showed that the ability of the 601 sequence to strongly position nucleosomes is lost in *S*.*cerevisiae* (Fig.2 b). We used our nucleosome density predictor [11] on the 601 sequence arrays used in this previous study and questioned whether it is able to predict the nucleosome density on a completely exogenous sequence, namely the Widom-601 sequence. In agreement with the experimental nucleosome densities, the predictor anticipates that the Widom-601 sequence is unable to position nucleosomes *in vivo* (Fig.2 b). The predicted density is relatively flat over the 167 bp long sequence, showing no evidence of nucleosome localization on the first 147 bp where the Widom-601 sequence is placed. Moreover, for both the 197 bp and the 237 bp long sequence, the predictor anticipated that the preferential position of the nucleosomes is around the linker, in agreement with the experimental results (Fig.2 c). The ability of our model to predict correctly the position of nucleosomes on exogenous sequences prompted to use it for a design purpose.

**Figure 2:**
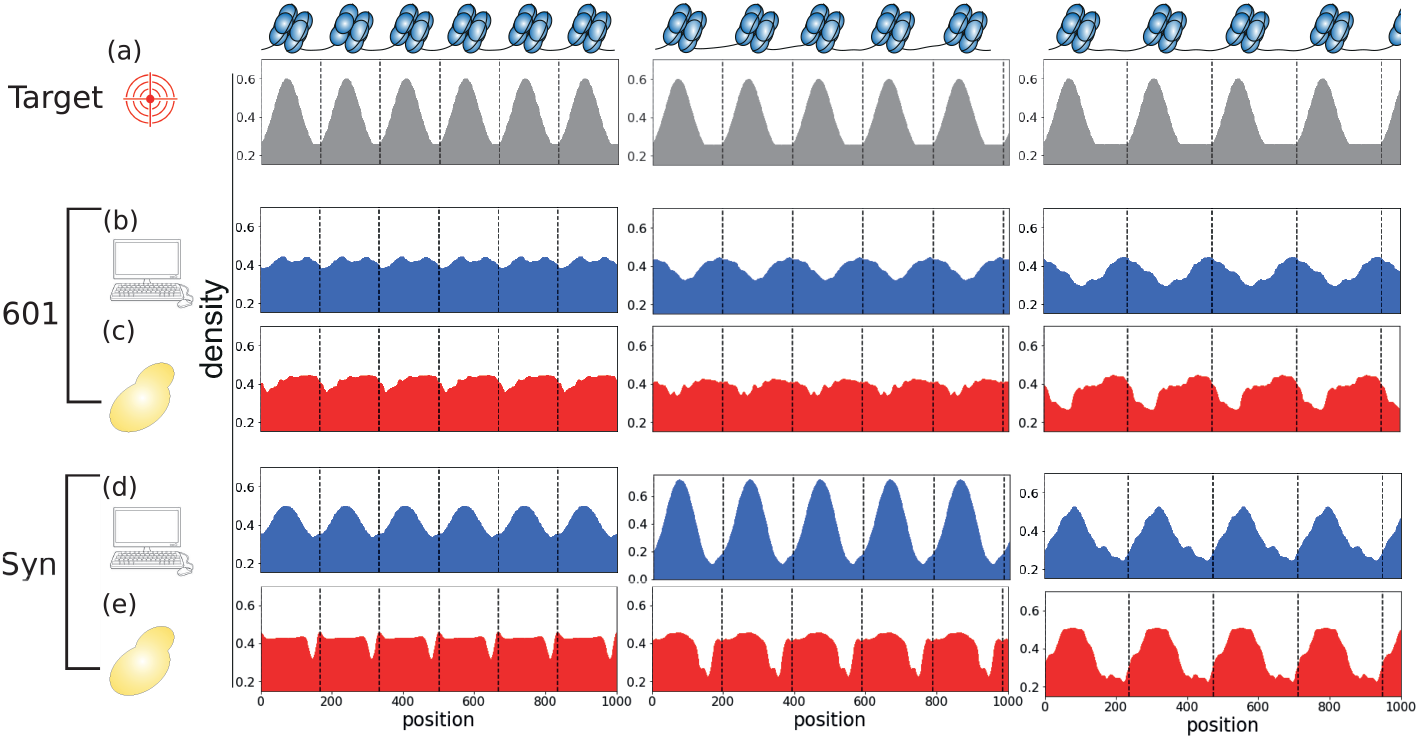
Target, predicted and experimental nucleosome densities on a 1000 bp long subset of Widom-601 repeats and of synthetic sequence repeats. (a) Target nucleosome densities corresponding (from left to right) to one nucleosome every 167 bp, 197 bp and 237 bp. (b,c) Normalized predicted (b) and experimental (c) nucleosome densitiy on a 1000 bp region of 601 repeats. (d,e) Normalized predicted (d) and experimental (e) nucleosome density on the 1000 bp subset of our synthetic sequence repeats. Densities are normalized so that the area under the curve is equal to the area under the curve of the target densities

### 3.2 Optimization of the monomer sequences for regular nucleosome positioning in tandem arrays

Starting from a random sequence, we computed the effect that would have on the predicted density all possible single mutations (Fig.1). We then selected one of these mutations in order to obtain a nucleosomal density corresponding to one nucleosome precisely occupying the first 147 bp and being excluded from the rest of the linker region. We devised an energy function (Fig.1 b) based on the distance between the predicted nucleosomal density on the tandem array sequence and a target density (in grey on Fig.1c). The k-MC methodology optimisation process typically converged after 20 to 50 steps, starting from a random DNA sequence (Methods and Fig.1). The distance to the target was rapidly divided by a factor 3 during the 5 first mutations steps, then slowly converged and finally stabilised around a value of 0.6. The predicted nucleosome densities corresponding to three time points sampled during the optimisation process illustrate the convergence of the predicted density towards the target function (Fig.1 c, right panels).

Quite surprisingly, for all initial random sequences we used, 5 to 20 changes out of 167 197 or 237 bp are always sufficient to provide a strong nucleosome positioning sequence (**Supplementary figure S2**). Starting from random DNA seeds and after many round of sequence optimisation, we could produce hundreds of positioning sequences with a predicted nucleosome at the desired dyad position (**Supplementary figure S3**). Three sequences corresponding to the three NRLs we wished to impose on our tandem arrays were selected for further study.

### 3.3 Experimental validation of the positioning capabilities of the synthetic sequences

In order to compare the positioning efficiency of our three *in silico* optimized sequences with the *in vitro* optimized Widom’s 601 sequence, we performed MNase-seq experiments on three *S*.*cerevisiae* strains in which repeated arrays containing 167 bp, 197 bp or 237 bp long monomers were engineered into the non essential YMR262 gene in chromosome XIII (**Supplementary Figure S4**, Online Methods and [9]). We obtained mono-nucleosomal fragments both in the genomic sequences (**Supplementary figure S5a, b, c**) and in the synthetic repeated regions (**Supplementary Figure S5d, e, f**). The experimental nucleosomal densities over the arrays as well as the corresponding predicted densities are shown on Fig.2. Our results unambiguously show that our sequence optimisation framework enables the generation of non-natural sequences that are able to produce such arrays for 167, 197 and 237 bp (Fig.2 d,e). For the 167 and 197 bp repeat, we observed a global preferential positioning of the nucleosome in the first 147 bp of the synthetic sequence with a flat peak which indicates that the nucleosome dyads (i.e. the fragments mid-points) are sharply distributed around the center of the first 147 bp (**Supplementary Figure S6a**). For the 197 bp long synthetic sequence, we also find a nucleosome well positioned on the first 147 bp of the sequence and a precisely positioned dyad (**Supplementary Figure S6b**). For the 237 bp long synthetic sequence tandem repeats, the experimental density exhibited a peak on the first 150 bp followed by a constant low density over 90 bp. Indicating that most of the nucleosomes were correctly positioned on the sequence. The central position of sequenced fragments distribution (**Supplementary Figure S6c**) also showed a strong enrichment in the first 147 bp although with a broader distribution, potentially resulting from several nucleosome preferred positions. The linker regions were as expected strongly depleted in nucleosomes.

### 3.4 Analysing the optimisation process of the synthetic sequences

In order to better understand how the sequence was designed to position the nucleosomes we made a quantitative analysis of the mutations that arise during the optimisation process. We first identified which regions of the sequence are the most important to design a positioning sequence by performing 1000 k-MCMC independent, 20 steps long, optimisations and recording at each step the energy changes associated with each of the possible mutation. The more important a given mutation is, the more is the associated change in energy, either positively or negatively. The absolute value of the average Δ*E* along the sequence suggests that the most important regions for nucleosome positioning are the linker region (in blue in Fig.3 a) and the 50 bp region surrounding the dyad axis of the nucleosome (in red in Fig.3). In order to go further into the sequence rules that define nucleosomes positions, we studied the motifs corresponding to the most important mutations (Online Methods). We split mutations into four groups: positioned either in the linker or in the dyad region and either increasing or decreasing the energy term. We then searched for typical 5 bp long DNA motifs within each group using STREME [18]. Two types of mutations could be extracted: mutations that construct one of the two known nucleosome repelling motives (poly(dA-dT) and poly(dCG)) and mutations that destruct these motives [16]. The sign of the resulting energy change is dictated by the position of the nucleotide within the monomer sequence: in the linker region, destructive mutations increase the energy while constructive mutations decrease the energy. In the dyad region the opposite occurs. These results are consistent with previous studies suggesting that nucleosome attractive sequences do not exist in *S*.*cerevisiae* genome and explain the inefficiency of the 601 sequence to position nucleosomes *in vivo* [5, 16].

**Figure 3:**
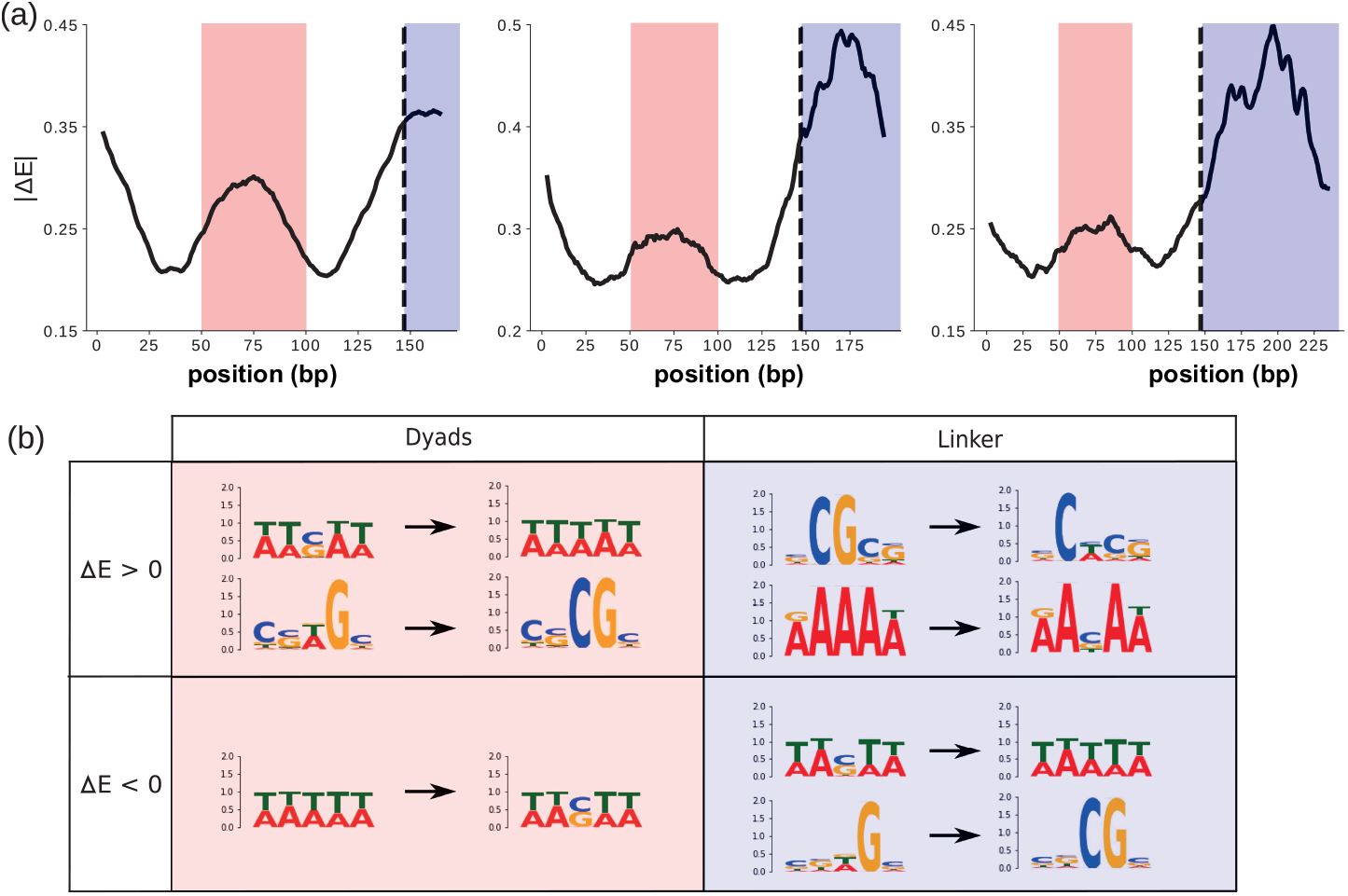
Positions on the sequence and typical motifs around important mutations during the optimisation process. (a) Average absolute change in energy due to mutations at each position along the 167 bp, 197 bp and 237 bp monomer sequence (from left to right). (b) DNA motifs enriched around important mutations before and after the mutation (respectively on the left and right of the black arrows).

## 4 CONCLUSION

In this study, we used the k-MC heuristic combined with a deep learning nucleosome density predictor to design an array of nucleosome positioning sequence. By assembling DNA tandem repeats of those synthetic sequence in *S*.*cerevisiae* genome, we validated that they were able to position a nucleosome on their first 147 bp and lead to the formation of regular arrays of various NRLs, including non-natural ones. The experimental control of the local NRL will enable the study of the structure of the nucleosomal fiber *in vivo* to decipher the causal link between nucleosome positioning, epigenetics marks, transcription or other related processes linked to chromatin structure. Our method, which combines a deep learning predictive method and the k-MCMC methodology, can be adapted to design DNA sequences with other designer characteristics and we expect it to play a growing role in the emerging field of synthetic genomics.

## 5 ACKNOWLEDGEMENTS

We thank members of the Structure and Instabilités des génomes and LPTMC laboratories for constructive comments during the realization of this work. Work was supported by core funding from CNRS, INSERM and MNHN. ALJ and ER were supported by a doctoral fellowship from the french ministry for Education, Research and Technology. EP was funded by grant ANR-15-CE11-0023-03 (HiResBac).

## 6 Availability of data and materials

Synthetic strains developed in this work are available upon request to JBB or AJ.

Raw sequencing reads for Mnase-Seq experimentsare available at the following address: https://dataview.ncbi.nlm.nih.gov/object/ PRJNA863754?reviewer=c8p620eemjlo6r3dvv6kcjoip

The code developed in this work is available on github at the following address: https://github.com/etirouthier/nuc_sequence_design

