## Supplementary Figures for "*In silico* design of DNA sequences for *in vivo* nucleosome positioning"

Table 1: **Synthetic sequences used for *in vivo* nucleosome positioning**

| repeat ID | sequence |
|---|---|
| 167-SSNP | ATAGAAGCTAAATAAGTAGTATGAACTGATAAGGGACACAAGAGAACGAAGAACATTTCCT CCGCAACTTCGAATGCTGGTGTGTCTCTAGCGACGTTGGTCTAATGTGTTTGTACTTTTTT GGTCTTACGATGCTAGTATTGAGTTATAATTAATCGCCCAACAAA |
| 197-2-SSNP | GAAGACTCTTCCGAGATAAGTTTTTTCCTTGATTGATAGATGAGTACGTCTGACCTATCTA GACCATGTGACTTCCGAGAGTGCTACATAGCTTCTACACATTCATCTATGAATCCTCAAAT GGAAAGGCTGTATAAAAATGCCTTCGCGTGAATGTTTAGGCGTAAGGAGATTCGGTTGGCG CGCTCCGTGAAACA |
| 237-SSNP | GATTTACTTTAGAGCTAAACTATATATAGTGATAATTAATGTTGCAGCGAGGGCGTCCACG TATCCGAGCGACACCAGTGGGCACCGACTCTACTTCTCCCGTTCCCGTCAACGCTAATATC TTCATATTTCAAACCTATCTTTTAATGTATAAATCCCTTAGGCCCCGCCCTAATGCAGGTG GATCGAAAAAGCTTAAAATTCTGGCTTTGAGTAAAGGTTATAATGTAATTTTAA |

Table 2: **Strains used in this study**

| Strain name | Genotype | Reference |
|---|---|---|
| YPH499 | MATa ura3-52 lys2-801_amber ade2-101_ochre trp1-$\Delta 63$ his3-$\Delta 200$ leu2$\Delta 1$ | SGD |
| ALY0 | YPH499 + pAL30 | [1] |
| AJ_Y2 | ALY0 ymr262-$\Delta$79::SSNP167 | This study |
| AJ_Y4 | ALY0 ymr262-$\Delta$79::SSNP197-2 | This study |
| AJ_Y5 | ALY0 ymr262-$\Delta$79::SSNP237 | This study |

Table 3: **Plasmid used in this study**

| Plasmid name | Description | Reference | plasmids/oligos |
|---|---|---|---|
| AJ_P1 | pRS413-TEF1p-Cas9-His | [1] | |
| AJ_P2 | pRS426-SNR52p-gRNA.YMR262-SUP34t | [1] | |
| AJ_P3 | pRS426-SNR52p-gRNA.SSNPfor-SUP34t | this study | AJ_P2 /O-33/O-34/O-35/O-36 |
| AJ_P4 | pRS425-Trp-SNR52p-gRNA.ARG4-SUP34t | this study | AJ_P2 /O-33/O-34/O-35/O-36 |

Table 4: **Oligos used in this study**

| Name | Sequence | Description |
|---|---|---|
| O-1 | AAGCGACGATAATAGTCATTGAGGTTG | fwd-Left-YMR/SSNP |
| O-2 | TTATCAGTTCATACTACTTATTTAGCTTCTATATTAATTAATCCCCTTCAGCACGCAGC | rev-left-YMR/SSNP-167 |
| O-4 | ATCAAGGAAAAAACTTATCTCGGAAGAGTCTTCATTAATTAATCCCCTTCAGCACGCAGC | rev-left-YMR/SSNP-197-2 |
| O-5 | ATCACTATATATAGTTTAGCTCTAAAGTAAATCATTAATTAATCCCCTTCAGCACGCAG | rev-left-YMR/SSNP-237 |
| O-6 | ATAATTAATCGCCCAACAAAACAAGGTTTCTCACTATCAAGATGTACTGG | fwd-right-SSNP-167-YMR |
| O-8 | CGTAAGGAGATTCGGTTGGCGCGCTCCGTGAAACAACAAGGTTTCTCACTATCAAGATGTACTGG | fwd-right-SSNP-197-2-YMR |
| O-9 | TCTGGCTTTGAGTAAAGGTTATAATGTAATTTTAAACAAGGTTTCTCACTATCAAGATG | fwd-right-SSNP-237-YMR |
| O-10 | CCCGTGGCACTTTACATCGTG | fwd-right-SSNP-YMR |
| O-11 | ATAGAAGCTAAATAAGTAGTATGAACTGATAAGGGACACAAGAGAACGAAGAACATTTCCTCCG | F1-SSNP-167 |
| O-12 | CCAACGTCGCTAGAGACACACCAGCATTCGAAGTTGCGGAGGAAATGTTCTTCGTTCTC | R2-SSNP-167 |
| O-13 | TGTCTCTAGCGACGTTGGTCTAATGTGTTTGTACTTTTTTGGTCTTACGATGCTAGTATTG | F3-SSNP-167 |
| O-14 | CATACTACTTATTTAGCTTCTATTTTGTTGGGCGATTAATTATAACTCAATACTAGCATCGTAAGA | R4-SSNP-167 |
| O-20 | ATCCATACTGTGGTTTTTCTTCTCAAAAAAGAACCTGGCAACTCATGCCGGA | R6-SSNP-197-2 |
| O-21 | GAAGACTCTTCCGAGATAAGTTTTTTCCTTGATTGATAGATGAGTACGTCTGAC | F1-SSNP-197-2 |
| O-22 | GCACTCTCGGAAGTCACATGGTCTAGATAGGTCAGACGTACTCATCTATCAATC | R2-SSNP-197-2 |

Continued on next page

Table 4: **Oligos used in this study**

| Name | Sequence | Description |
|---|---|---|
| O-23 | GACCATGTGACTTCCGAGAGTGCTACATAGCTTCTACACATTCATCTATGAATCCTCAAAT | F3-SSNP-197-2 |
| O-24 | TTCACGCGAAGGCATTTTTATACAGCCTTTCCATTTGAGGATTCATAGATGAATGTG | R4-SSNP-197-2 |
| O-25 | GTATAAAAATGCCTTCGCGTGAATGTTTAGGCGTAAGGAGATTCGGTTGGCGCG | F5-SSNP-197-2 |
| O-26 | CTTATCTCGGAAGAGTCTTCTGTTTCACGGAGCGCGCCAACCGAATCTCCTTACGCC | R6-SSNP-197-2 |
| O-27 | GATTTACTTTAGAGCTAAACTATATATAGTGATAATTAATGTTGCAGCGAGGGCGTCCA | F1-SSNP-237 |
| O-28 | GAGAAGTAGAGTCGGTGCCCACTGGTGTCGCTCGGATACGTGGACGCCCTCGCTGCAAC | R2-SSNP-237 |
| O-29 | GGCACCGACTCTACTTCTCCCGTTCCCGTCAACGCTAATATCTTCATATTTCAAACCTATC | F3-SSNP-237 |
| O-30 | GCATTAGGGCGGGGCCTAAGGGATTTATACATTAAAAGATAGGTTTGAAATATGAAGAT | R4-SSNP-237 |
| O-31 | CTTAGGCCCCGCCCTAATGCAGGTGGATCGAAAAAGCTTAAAATTCTGGCTTTGAGTAA | F5-SSNP-237 |
| O-32 | TATAGTTTAGCTCTAAAGTAAATCTTAAAATTACATTATAACCTTTACTCAAAGCCAGAATTTTAAG | R6-SSNP-237 |
| O-33 | GAGCAGCTGCAGTATTTGAATGCAC | Southern_probe_for |
| O-34 | GCTACCCGCACCGTATACAAAAAGG | Southern_probe_rev |

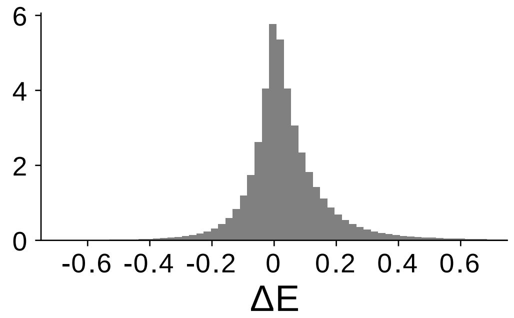


Figure S1: **Distribution of energy changes associated with all the mutations obtained during 1000 optimisation processes of 20 steps each.** The y axis has a log10 scale.

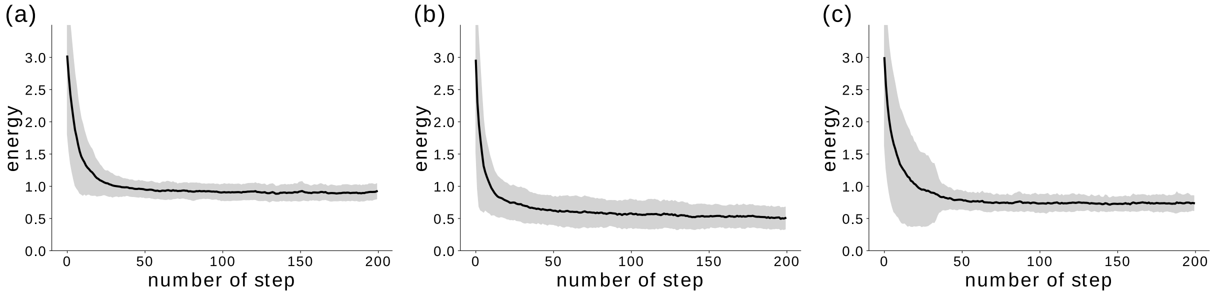


Figure S2: **Evolution of the energy during the k-MC procedure.** (a,b,c) Evolution of the mean energy (dark line) and the standard deviation (grey area) during the k-MC procedure for synthetic sequences of respectively (167 a, 197 b and 237 c) bp long. The mean energy and the standard deviation are obtained using 100 synthetic sequences.

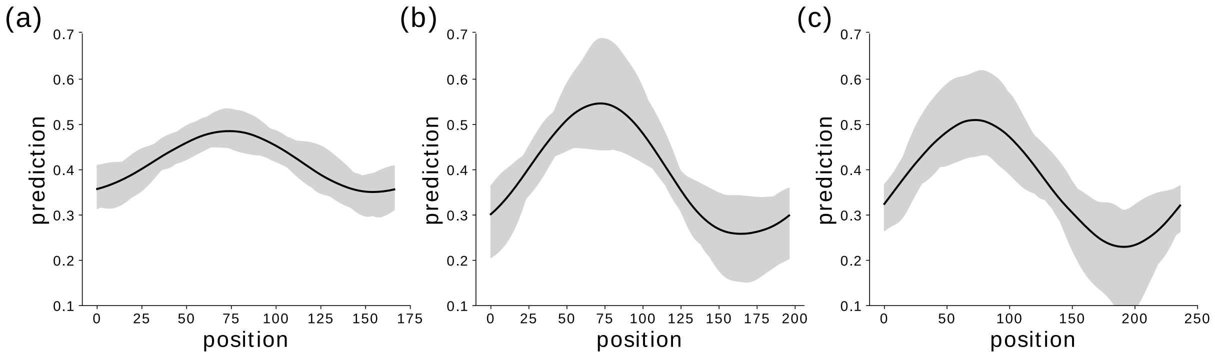


Figure S3: **Maximal, average and minimal predicted nucleosome densities for a set of 100 examples of optimized predicted densities.** The area between the minimal and the maximal predicted nucleosome density at each base pair of synthetic sequence (167 bp (a), 197 bp (b), 237 bp (c)) is filled in grey. The mean predicted nucleosome density is represented as a the dark line.

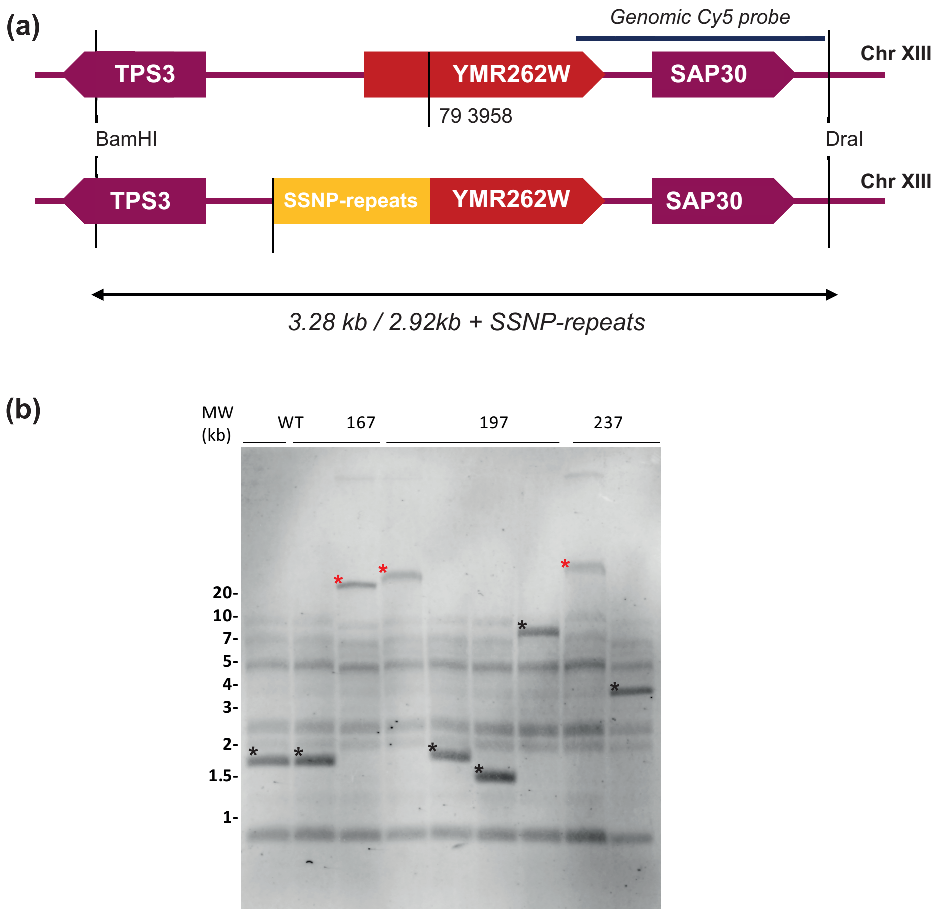


Figure S4: **Oligonucleotide assembly at the *YMR262W* locus.** (a) Diagram of the assembly region before and after Cas9-assisted recombinational assembly. (b) Southern Blot analysis of yeast recombinant strains. Genomic DNA from strains containing SSNP repeats were digested with enzymes BamHI and DraI. Samples were electrophoresed, blotted and hybridized with a Cy5 genomic probe. Bands corresponding to tandem DNA repeats (showing variable size compared to the control lane) are marked by an asterix. 167, 197, and 237 design respectively the size of the monomer assembled as tandem DNA repeats upon recombination in vivo. Strains carryong repeats estimated at >20kb were kept for further analysis (red star).

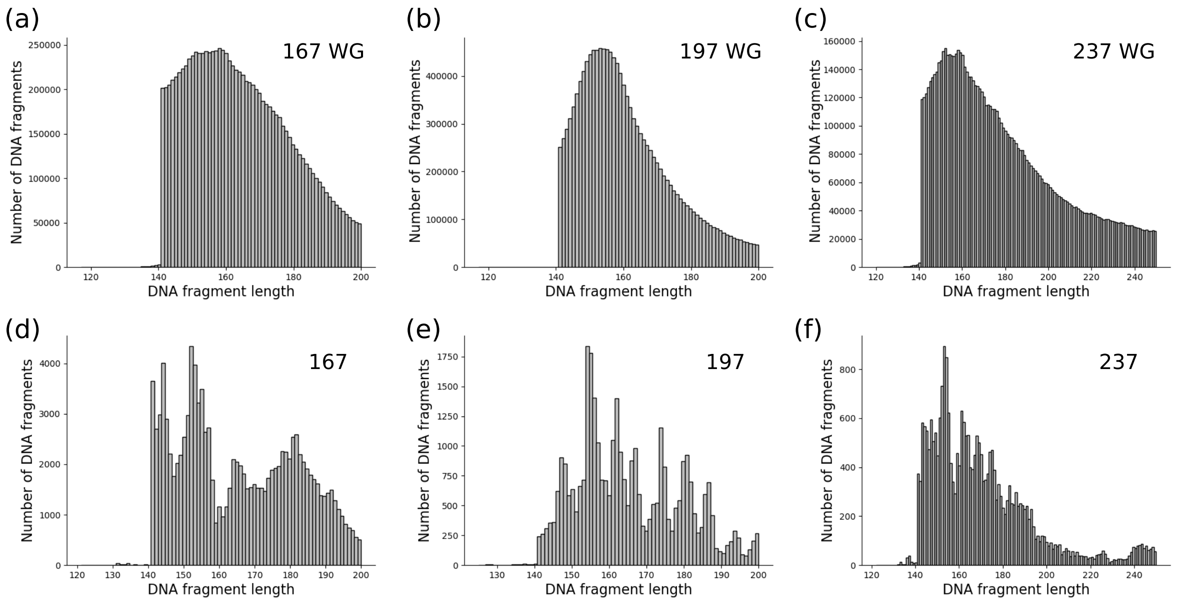


Figure S5: **Length distribution of mononucleosomal DNA samples genome wide and in the repeated synthetic region.** Histogram of the lengths of all nucleosome fragments obtained for a typical sample considering fragments from whole genome for each of the three 167 bp (a), 197 bp (b) and 237 bp (c) strains and separately considering only fragments overlapping the synthetic repeated region ((d) 167 bp, (e) 197 bp, (f) 237 bp.

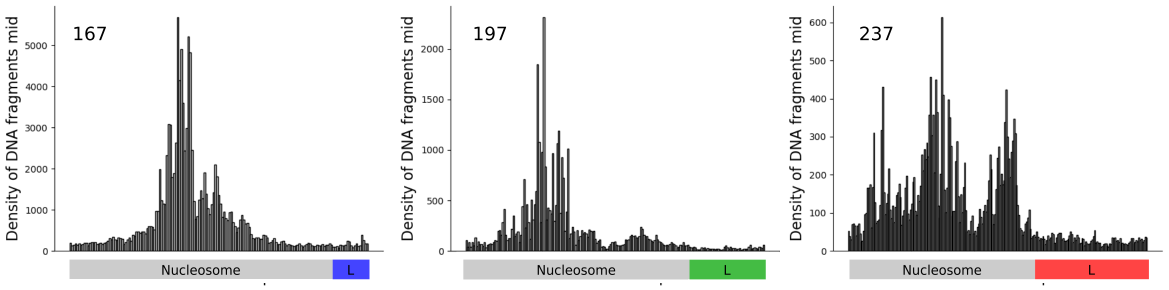


Figure S6: **Centre of sequenced fragments positions on synthetic repeated region.** Density profiles for synthetic sequence of 167 bp long (a), 197 bp (b) and 237 (c). The grey rectangle correspond to the first 147 bp where we aim to position a nucleosome and the blue, green and red rectangle respectively correspond to the 20 bp, 50 bp and 90 bp long linker.
